# Paratope Prediction using Convolutional and Recurrent Neural Networks

**DOI:** 10.1101/185488

**Authors:** Edgar Liberis, Petar Veličković, Pietro Sormanni, Michele Vendruscolo, Pietro Liò

## Abstract

Antibodies play an essential role in the immune system of vertebrates and are vital tools in research and diagnostics. While hypervariable regions of antibodies, which are responsible for binding, can be readily identified from their amino acid sequence, it remains challenging to accurately pinpoint which amino acids will be in contact with the antigen (the paratope). In this work, we present a sequence-based probabilistic machine learning algorithm for paratope prediction, named Parapred. Parapred uses a deep-learning architecture to leverage features from both local residue neighbourhoods and across the entire sequence. The method outperforms the current state-of-the-art methodology, and only requires a stretch of amino acid sequence corresponding to a hypervariable region as an input, without any information about the antigen. We further show that our predictions can be used to improve both speed and accuracy of a rigid docking algorithm. The Parapred method is freely available at https://github.com/eliberis/parapred for download.

## I. INTRODUCTION

Antibodies are a special class of proteins produced by the immune system of vertebrates to neutralize pathogens, such as bacteria or viruses. They act by binding tightly to a unique molecule of the foreign agent, called the antigen. Antibody binding can mark it for future destruction by the immune system or, in some instances, neutralize it directly (*e.g*. by blocking a part of a virus essential for cell invasion). Typical antibodies are tetrameric—made of two immunoglobulin (Ig) heavy chains and two Ig light chains—and have a Y-shaped structure, where each of the two identical tips contains a binding site (paratope). The base of the Y mediates the ability of an antibody to communicate with other components of the immune system.

The paratope is typically contained within the hyper-variable regions of the antibody which are also referred to as *complementarity determining regions* (*CDRs*). In the structure of an antibody, CDRs are located within binding loops, three on each heavy chain (H1, H2, H3) and three on each light chain (L1, L2, L3). The variability of the CDR sequences allows antibodies to form complexes with virtually any antigen. This binding malleability of antibodies is increasingly harnessed by the biotechnological and biopharmaceutical industry; indeed, monoclonal antibodies are currently the fastest growing class of therapeutics on the market (Ecker *et al.* [1], Reichert [2]).

Novel antibodies that bind a target of interest can be obtained using well-established methods based on animal immunisation or on in vitro technologies for screening large laboratory-constructed libraries (Leavy [3]). However, for applications in research, diagnostics, and therapeutics, some degree of engineering is required to optimise certain properties, such as binding affinity, stability, solubility, or expression yield (Chiu *et al.* [4]). Rational engineering decisions become easier if detailed knowledge about an antibody under scrutiny is obtained (Chiu *et al.* [4], Sormanni *et al.* [5]). However, especially at the early stages of an antibody discovery campaign, only the sequence and an estimate of the binding affinity are usually available. Therefore, computational methods that can accurately predict molecular traits using just the amino acid sequence have a great potential for accelerating antibody discovery by assisting lead selection or facilitating property engineering.

For instance, hypervariable regions contain 40–50 amino acid residues, whereas typically less than 20 actually participate in binding (Esmaielbeiki *et al.* [6]), and some may even fall outside of the traditional definition of the CDRs (Kunik *et al.* [7]). The ability to accurately map the paratope would enable to pinpoint residues that are involved in binding, leaving others as candidate mutation sites that can be exploited to optimise other molecular traits, such as solubility or stability, without compromising the binding activity. In addition, as we show in this work, accurate paratope prediction can improve accuracy and speed of docking simulations, making structural models more reliable and easier to obtain.

In this work, we introduce the *Parapred* method for sequence-based prediction of paratope residues. Parapred improves on earlier methods for paratope prediction (Krawczyk *et al.* [9], Kunik *et al.* [7], Olimpieri *et al.* [8], Peng *et al.* [10], Tsuchiya & Mizuguchi [11]) by using deep learning methods and larger antibody datasets. Our method only requires the amino acid sequence of a CDR and four adjacent residues as its input, which, in contrast to structural data, can be readily obtained experimentally. For simplicity, we only consider antigens that are themselves proteins, which are the vast majority of known antibody targets.

“Deep learning” specifically refers to the process of building machine learning models consisting of multiple layers of non-linear operations, where each successive layer automatically learns more abstract representations (*features*) of the data using the features extracted by the previous layer (Goodfellow *et al.* [12, p. 1]). A key advantage of deep learning over traditional machine learning methods is that it can perform automated feature extraction directly from raw input data, thus eliminating the need for a domain expert to manually engineer features (Goodfellow *et al.* [12, p. 4]). Automatically learned features are often found to be superior to manually engineered ones, contributing to the widespread success of deep learning in a range of fields. In particular, Parapred builds upon convolutional and recurrent neural networks, which achieved state-of-the-art results in object recognition (Krizhevsky *et al.* [13]) and machine translation (Wu *et al.* [14]) tasks, among others. Deep learning has already been successfully applied to address problems in protein science, including the prediction of structure (Li *et al.* [15]), function (Tavanaei *et al.* [16]) or binding sites (Alipanahi *et al.* [17]). To the best of our knowledge, this work is the first application of modern deep learning to antibody-antigen interactions.

## II. METHOD

### A. Data acquisition and preprocessing

To train and test our models, we used a subset of the Structural Antibody Database (SAbDab) (Dunbar *et al.* [18]), which contains antibody and antigen crystal structures. Entries in SAbDab were filtered (using the SAbDab web interface on 27 June 2017) to obtain a non-redundant set of antibody-antigen complexes with the following properties: (1) antibodies have both *V*_*H*_ and *V*_*L*_ domains present, (2) structure resolution is better than 3Å, (3) no two antibody sequences have > 95% sequence identity, (4) no two antigen sequences > 90% identical, and (5) each antibody has at least 5 residues in contact with the antigen. The final dataset contains 239 bound complexes (Supplementary Table C).

To construct the input, we identify the CDRs within the sequence of each antibody using the Chothia numbering scheme (Al-Lazikani *et al.* [19]). We augment the CDR sequences with two extra residues at both ends, as these residues are also known to sometimes engage in binding (Krawczyk *et al.* [9], Kunik *et al.* [7]). These extended CDR sequences are the input of the Parapred method and are processed individually.

Amino acid sequences have to be encoded as tensors prior to being processed by the model (Figure 1):

- Each amino acid sequence is encoded as a ‘row’ in a 3D matrix. As CDR sequences are usually of different length, they are padded with zero vectors to the length of the longest sequence. This is necessary for fast batch tensor processing provided by deep learning frameworks.
- Each element in a matrix encodes an amino acid residue and is itself a vector consisting of two concatenated parts:

– *One-hot encoding* of the type of the residue (20 possible amino acid types + 1 extra, representing an unknown type). The type is encoded using a 21-dimensional vector, where all elements are set to 0 and one element, corresponding to the actual type of the amino acid, is set to 1.
– Seven additional features, summarised by Meiler *et al.* [20], which represent physical, chemical and structural properties of each type of amino acid residue (Supplementary Table A).

**Fig. 1.**
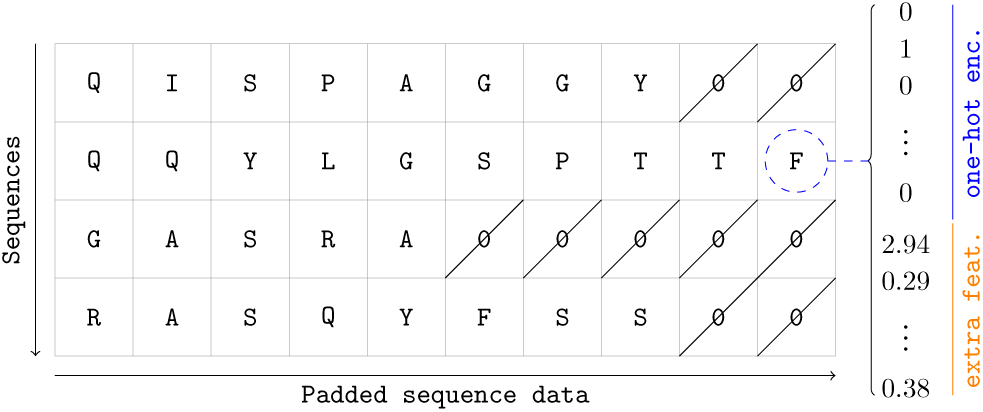
An example of encoded amino acid sequences. An amino acid residue is represented by a feature vector which consists of one-hot encoding and some extra features. To cope with different sequence lengths, each sequence is padded to the length of the longest one.

The final dataset contains 1434 sequences for the algorithm to learn from (239 antibody/antigen complexes × 6 CDRs each).

### B. Building a deep learning model

The paratope prediction problem can be formalised as a binary classification problem between two classes of residues: those that do not participate in binding (Class 0) and those that do (Class 1). Following previous conventions (Krawczyk *et al.* [9]), we define binding residues as those with at least one atom found within 4.5Å of any of the antigen atoms. Thus, for each residue in a sequence, the algorithm will output the probability *p* of it being in class 1 versus being in class 0 (the likelihood of binding).

Our model uses six prominent architectural developments in deep learning:

*1) Multilayer perceptrons (MLP):* Neural networks can be thought of as a set of interconnected units, called *neurons* or perceptrons, each of which performs a simple computation.

Neurons are typically arranged in *layers*, where each neuron in a layer is connected to the output of every neuron in the previous layer. A layer with this kind of connection is called *fully-connected*. The neural network itself is constructed as a series of such layers—the data is transformed in turn by every layer as it flows through the network. This architecture is known as a *deep feed-forward neural network* or a *multilayer perceptron* (MLP).

Neural network architectures are extensively used for machine learning tasks that can be reformulated as function approximation problems. We would like a network to learn to approximate some target function *ƒ*: *X* → *Y* using a set of known input / output pairs for it (*supervised learning* setup). For example, for paratope prediction, *x* ∈ *X* could be a vector encoding a residue and *y* ∈ *Y* = {0,1} could indicate whether the residue participates in binding.

The signals between neurons are real numbers and the neuron computes its output as follows:

- A neuron computes a weighted sum of its inputs (x) and adds a constant term to it. The coefficients by which every input is scaled are called *weights* (**W**) and the constant term is called the bias (*b*). The weights and bias constitute a set of adjustable *parameters* of a neuron.
- Some non-linear *activation function σ* is applied to the sum to produce the output. The activation function introduces a non-linearity necessary to model complex functions.

We can compactly write the transformation performed by all neurons in a layer as a single weight matrix multiplication and bias vector addition:

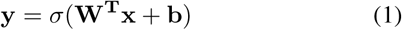

*2) Recurrent Neural Networks (RNN):* We can design a neural network which processes every element in a sequence in turn. The key idea behind RNNs is to iteratively apply a simple processing block, called RNN *cell*, to obtain a summarised representation of a sequence up to any point. Figure 2 shows a computation graph of an RNN—the cell iteratively consumes inputs (x) by computing a function of x and the previous state of the cell *s*.

We use the *Long Short-Term Memory (LSTM)* (Hochreiter & Schmidhuber [21]) cell which is able to learn long-range dependencies in sequences. The computation performed by an LSTM cell consists of the following steps:

- An LSTM cell holds the state *s* in two vectors: C (“memory”) and h (previous output). Input x and state vector h are concatenated before being processed in four steps:

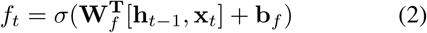

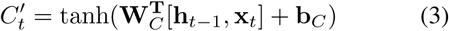

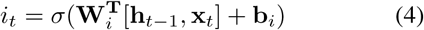

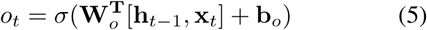 where tanh is the element-wise hyperbolic tangent and *σ* is the logistic sigmoid function 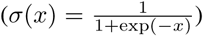. Matrices **W** and vectors b are parameters learned by the network.
- The new cell state *C*_*t*_ and *h*_*t*_, as well as the output *y*_*t*_ is given by:

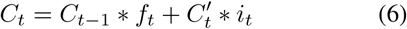

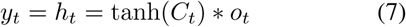

where _*_ is the element-wise vector multiplication.

Capturing dependencies between an output and later inputs is necessary for amino acid sequences because they don’t have a canonical direction (reading a sequence left to right is equivalent to reading it right to left). To achieve this, we use a *bidirectional RNN* (Schuster *et al.* [22]) which introduces a second pass going in the opposite direction (see Figure 2).

RNNs enable the model to capture features which span the entire input sequence.

*3) Convolutional Neural Networks (CNN):* Amino acid residues are known to interact with other residues and prefer some kinds of amino acids more than others as their neighbours (Xia *et al.* [23]). A paratope prediction model can exploit such preferences by processing every residue together with its neighbourhood to learn useful local patterns first and only then use an RNN to learn aggregate features of the entire sequence.

Spatially local features can be extracted using *convolutional layers*, typically found in *convolutional neural networks (CNNs)*.

A convolutional layer is similar to a single-layer MLP discussed previously, only it uses a convolution operation instead of matrix multiplication. A convolution operation for sequences is defined as:

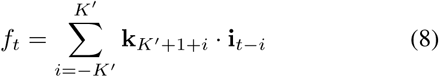

where **i**_*t*_ and *ƒ*_*t*_ are elements of the input and output sequences at position *t*, respectively, and **k** ∈ ℝ^*K* × *C*^ is a *kernel* of size *K* = 2*K*′ + 1 (w.l.o.g assume that the kernel has an odd number of elements; *C* refers to the dimensionality of the input). This computation is visualised in Figure 3.

The kernel is applied this way at every position of the input sequence to produce the output sequence. For positions where kernel spans beyond the input sequence, we assume the input is padded with zero vectors: **i**_*t*_ = 0 for *t* ≤ 0 or *t* > *T*. The input and kernel elements themselves are vectors with multiple *channels*—name comes from an analogy with images: each pixel in an image has 3 dimensions: red, green and blue channels—*e.g*. an encoded residue would have 28 dimensions / channels (20 + 1 amino acid type one-hot encoding + extra 7 features, as described earlier).

Convolution performs a weighted summation over all dimensions of input elements to produce a single number (the sum of vector dot products). The fact that the same small kernel is applied to every position in the input sequence allows it to detect input patterns regardless of their position. Learnable parameters of a convolutional layer are its kernels; multiple output channels can be produced by using several different kernels (*filters*).

*4) Residual Connections:* Residual connections (He *et al.* [24]) act as a shortcut connection between inputs and outputs of some part of a network by adding inputs to outputs. Such shortcut can be added around the convolutional feature extractor—if the local feature extractor is supposed to learn some function *h*(x), with the shortcut connection it only has to learn the residual *h*(x) - x which is often easier to optimise for. The shortcut also enables the rest of the model to learn both from original inputs and extracted local features, and acts as a complexity controller by effectively allowing the network to adjust its depth.

*5) Exponential linear units as activation functions:* Activation functions introduce a non-linearity which is necessary to model complex functions. Experimenting with the activation function’s behaviour can improve the training process. We use the *Exponential Linear Unit (ELU)* (Clevert *et al.* [25]) activation function which makes the network more robust to noise and faster to train. The function is given by:

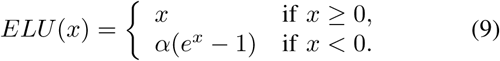

We use *α* = 1.

*6) Model regularisation:* Deep learning models often have to be regularised to prevent overfitting—a phenomenon where a network memorises training examples (and noise) instead of modelling the underlying relationship. We use two regularisation methods:

- *Dropout* (Srivastava *et al.* [26]) is a computationally efficient regularisation method. The main idea of Dropout is to discard some intermediate results of the network at every training iteration with a certain probability *p*. This discourages the network from learning to rely on a particular subset of inputs.
- *L*_2_ regularisation (aka *weight decay*) adds an extra term—an *L*_2_ norm of a layer’s weights—to network’s optimisation objective, which penalises weights if they grow too large during training.

**Fig. 2.**
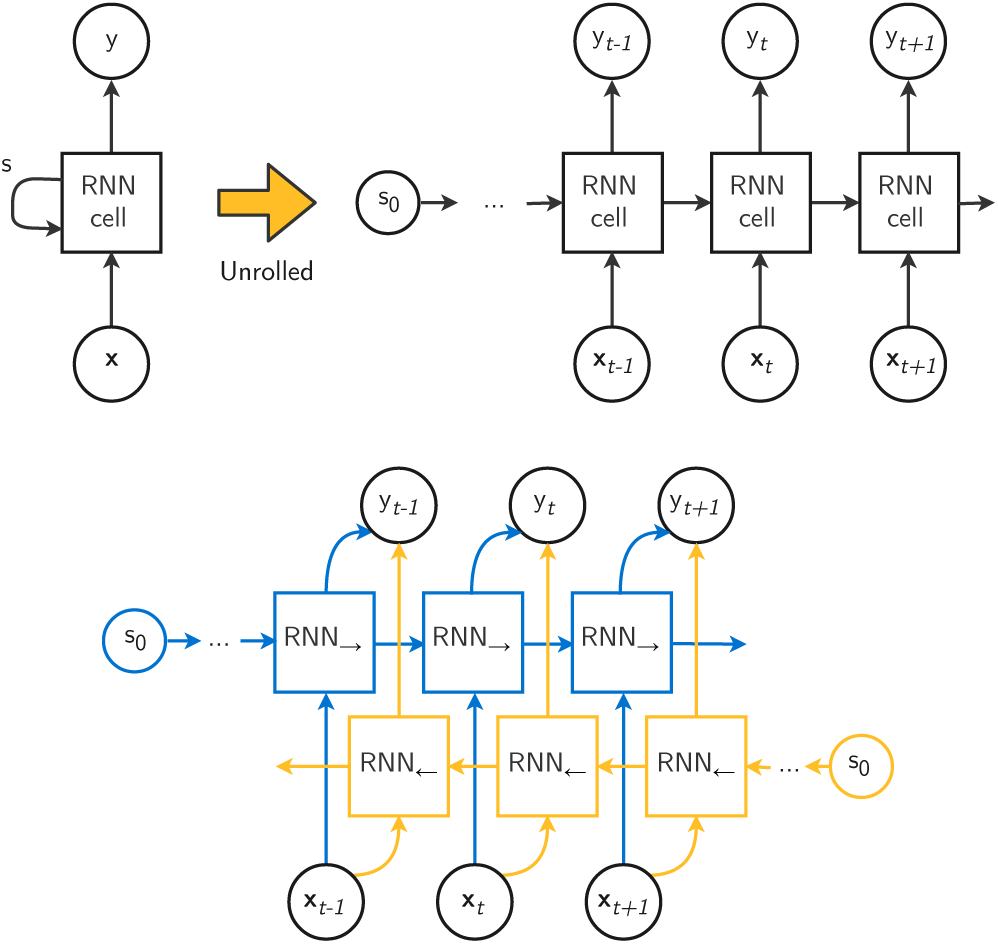
The computation graph of RNNs used in this work. **Top:** the computation graph before (left) and after (right) unrolling. The same RNN cell is used to process every element of the input sequence. **Bottom:** unrolled graph of a bidirectional RNN. The inputs are passed through two different RNN cells (one for each direction) and the network’s output at time *t* is an aggregation (here—concatenation) of the two cells’ outputs.

**Fig. 3.**
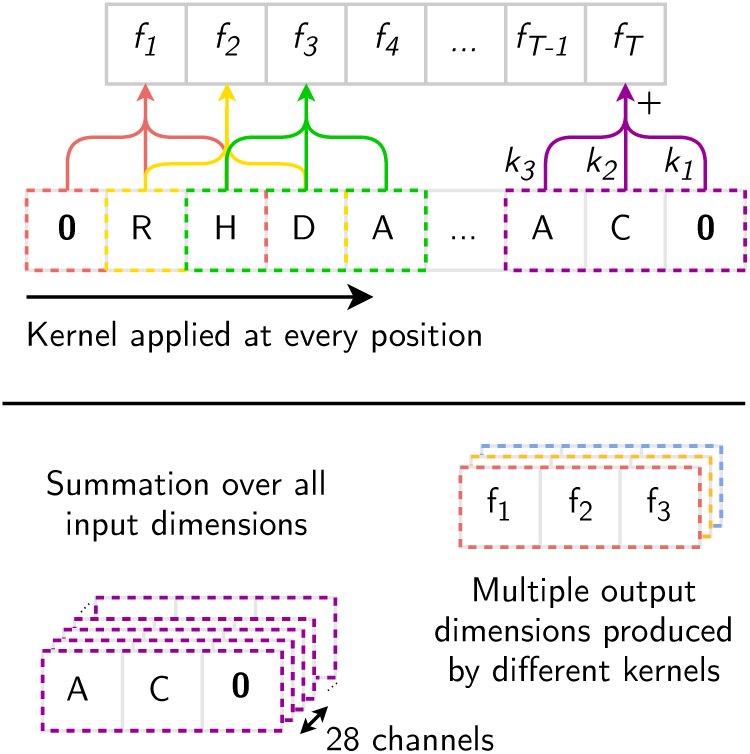
An example of 1D convolution with kernel size 3. Outputs are computed by applying a kernel at each position in the input sequence.

### C. Experimental setup

The software was developed in Python using TensorFlow deep learning framework (Abadi *et al.* [28]) and Keras API (Chollet *et al.* [27]). Overall, the network’s computation consists of the following steps (Figure 4):

1. Encoded sequences (CDRs with 2 extra residues) are processed by a convolutional layer (regularised with an *L*_2_ term scaled by 0.01) with 28 kernels, each spanning a neighbourhood of 3 residues. ELU activation is applied to the convolution results.
2. Residual connection is implemented by adding the original input sequences to the convolution output.
3. Resulting features are processed by a bidirectional LSTM with state size 256. The network applies Dropout with *p* = 0.15 to RNNs input and Dropout with *p* = 0.2 to RNNs recurrent connections.
4. Dropout with *p* = 0.3 is applied to the RNNs output and individual feature vectors are processed by a single-output fully-connected network with logistic sigmoid activation function (to bring the output to the range of probabilities). Network’s weights are regularised using an *L*_2_ term scaled by 0.01.

**Fig. 4.**
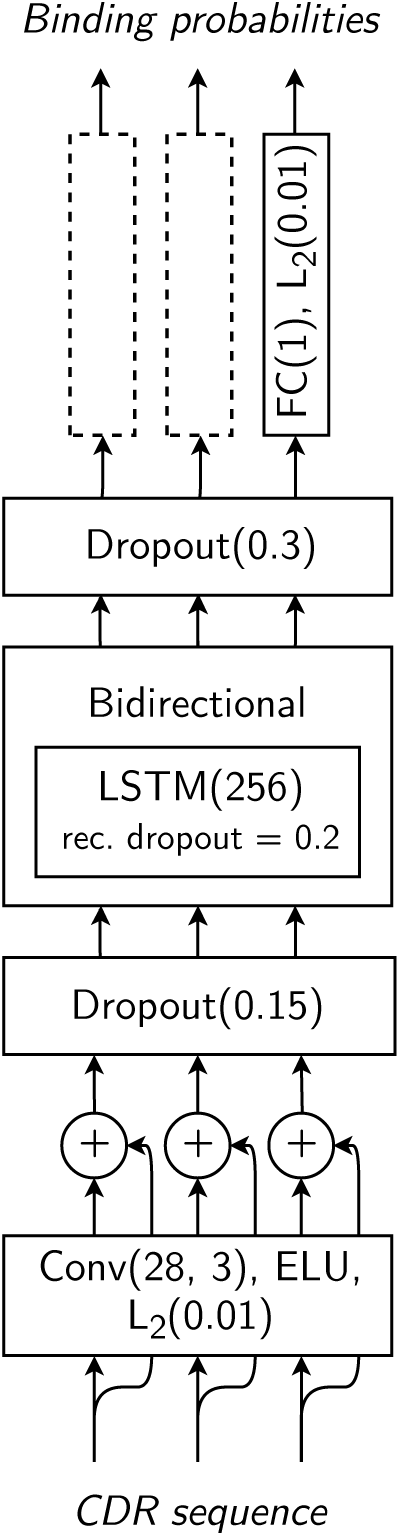
The architecture of our paratope prediction model.

The model’s architecture could be easily augmented with layers that are able to process the 3D structure of an antibody in conjunction with its amino acid sequence. However, such sophisticated architectures would require a much larger training dataset (at least l0x more 3D structures) which is not available at this time. Training this kind of model would also require a way of efficiently exploiting cross-modality during feature extraction (Veličković *et al.* [34]).

Neural network training is a function optimisation problem, where we aim to find a local or global optimum of the optimisation target (aka *loss*) with respect to network’s parameters. This should be a differentiable measure of how well the neural network approximates the target function. We use the *binary cross-entropy* loss, a popular choice for binary classification problems:

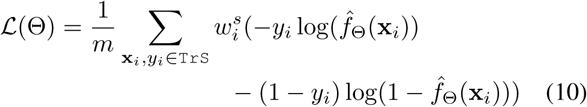

where TrS is the training set of size *m*, 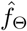 is the function computed by the network with parameters Θ and 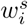 is the sample weight (described later).

To find a loss minima, we use the *Adam* (Kingma & Ba [29]) optimiser with base learning rate setting of 0.01 for the first 10 epochs and 0.001 otherwise. The network is trained with 32 samples at once (aka *batch size*) for 16 epochs (iterations over the entire training set).

The dataset has an uneven number of binding (positive) and non-binding (negative) residues—3.4x more negative samples. The cross-entropy loss function (Equation 10) equally penalises misclassified positive and negative samples, which allows the model to keep the overall loss low by preferring to predict that residues will not bind. This achieves good classification accuracy but hinders the model’s ability to learn to identify positive samples. This can be improved by penalising misclassified positive samples more—the per-sample loss is scaled by the *sample weight* 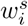 which we set to a 2.5x higher value for positive samples.

## III. RESULTS

### A. Model results

We used the *10-fold cross-validation* technique to assess the model performance on multiple dataset splits. This technique randomly partitions the data into ten subsets and trains the model ten times. Each time, a different subset is selected for testing, and the method is trained on the sequences belonging to the other nine. Because the results may vary between cross-validation runs due to the initial random partition of the data, the random initialisation of the network parameters, and Dropout, the 10-fold cross-validation is repeated ten times, which also enable to calculate confidence interval of the mean values of each performance indicator.

To measure the performance of the binary classifier we use a number of standard metrics:

- **Recall:** (Also known as *sensitivity* or *true positive rate*.) The proportion of positive samples classified correctly 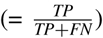.
- **Precision:** (Also known as *positive predictive value*.) The proportion of actual positive samples among all samples predicted to be positive 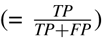.
- **F-score**: Precision and recall measure two independent properties of a classifier—F-score combines both of them by measuring their harmonic mean.
- **Matthews correlation coefficient (MCC):** MCC is a popular measure of binary classification quality for unbalanced datasets.
- **Precision-recall (PR) curve:** Aforementioned metrics require a particular classification threshold. However, the user can vary the threshold to adjust the confidence of results they get. For example, setting the threshold to a high value would label only very few residues as positive (those that the model is very certain about), so the classification would have low recall but high precision. Conversely, setting the threshold to a low value would yield high recall but low precision results. To visualise this relationship, we plot precision at different levels of recall obtained by varying the threshold.
- **Receiver operating characteristic (ROC) curve:** ROC curves show the relationship between the true positive rate (recall) and the false positive rate (the proportion of false positives among all negative samples, 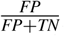), similarly obtained by varying the threshold. Area under the ROC curve (**ROC AUC**) is often used to compare classifiers.

Table 1 shows the F-score, MCC and ROC AUC performance metrics of Parapred. Narrow confidence intervals indicate consistent performance across cross-validation rounds. Furthermore, the results show that our model outperforms the current state of the art predictor, *proABC* (Olimpieri *et al.* [8]), without needing the entire antibody sequence or extra features such as the germline family or antigen volume.

**Table 1.** Performance indicators of the Parapred method with 95% confidence intervals (top row) and of the proABC method (bottom row). The F-score and MCC metrics use a classification threshold of 0.4913739, obtained by maximising Youden’s index (Youden [30]).

| Model | F-score | MCC | ROC AUC |
| --- | --- | --- | --- |
| *Parapred* | $0.705 \pm 0.003$ | $0.572 \pm 0.005$ | $0.883 \pm 0.002$ |
| *proABC* | — | 0.522 | 0.851 |

Figure 5 shows the precision/recall curve of the Parapred method (in blue) which indicates a substantial improvement over Antibody i-Patch (orange points). This improvement is particularly relevant given that, in contrast to Parapred, Antibody i-Patch requires a structure or a homology model of the antibody and the antigen it binds to.

**Fig. 5.**
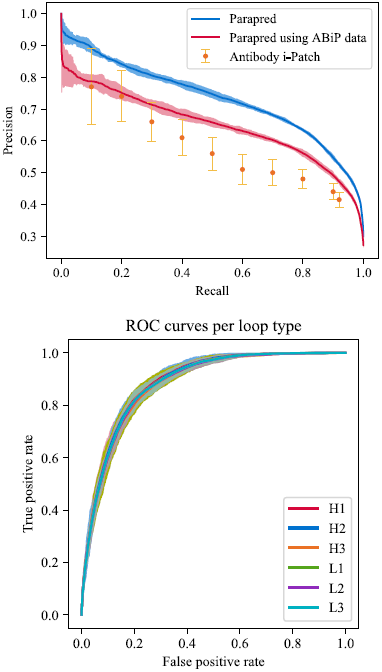
Parapred performance curves. **Top:** precision-recall curves of Parapred, obtained from 10-fold cross-validation runs, when trained on our dataset (blue) and Antibody i-Patch’s dataset (red), together with the PR values reported by the authors of Antibody i-Patch (orange). **Bottom:** ROC curves of Parapred, separated by loop type. Shaded areas / bars show 95% confidence bounds (2 standard errors).

We investigated to what extent the performance improvement originates from using a larger dataset (239 complexes vs. 148 of Antibody i-Patch) and to what from the deep-learning-based architecture of Parapred. To assess this, we measured Parapred’s performance when trained on the Antibody i-Patch’s dataset (red curve in Figure 5). We find that our method achieves significant precision improvements for recall values > 0.5, which is typically the most useful range. We conclude that the deep-learning-based architecture of Parapred is able to capture a richer set of features leading to better classification, even though it uses less explicit information about the antibody (Parapred does not require structural data or any information about the antigen it binds to). The leap in performance from the red to the blue curve is in agreement with the observation that deep models thrive in environments with a larger number of data points to learn from (Goodfellow *et al.* [12, p. 430]).

Our encoding of an amino acid sequence does not include information about the CDR loop type it originated from, so the model would not be able to capture loop type-specific features. Figure 5 shows the ROC curves of our model’s predictions, separated by CDR types. The graph shows that the model is able to predict binding residues equally well for all six CDR types despite lacking loop type information. We found that including the loop type made no appreciable difference to the performance.

### B. Docking improvements

We show the usefulness of Parapred by integrating its predictions with the PatchDock rigid protein docking algorithm (Duhovny *et al.* [31]).

PatchDock works with 2 protein molecules in the PDB format and searches for suitable orientations for one of the molecules—conventionally, the antigen—“onto” the other, such that the two form an antibody-antigen complex. The algorithm produces several hundred candidate orientations of the antigen, called *decoys*, which are ranked in the output by an internal scoring function. The algorithm also provides facility to guide the search process by pre-specifying potential binding site residues.

Decoys can be stratified into 4 quality classes—high (***), medium (**), low (*) or unclassed—based on how close the computed orientation of the antigen is to the true (*native*) orientation recorded in the dataset. The classification uses the CAPRI criteria (see Appendix B).

The usefulness of Parapred was measured by running PatchDock with three potential binding sites of the antibody molecule: (1) the CDRs, (2) the actual paratope and (3) binding residues predicted by Parapred (residues with > 0.67 binding probability to match the number of residues in the actual paratope). We picked 30 antibody-antigen complexes at random (highlighted in Appendix C) to be run through the docking algorithm and used the remaining 209 to train the model.

We recorded the best class decoy in the top 10 and top 200 decoys, as ranked by PatchDock, for each of the 30 structures. As a hint, we also supplied the antigen’s binding site as residues within 5Å of the real epitope. PatchDock was run with default parameters.

Docking results are shown in Table 2—supplying just the CDR gives the worst performance, however, supplying our model’s predictions achieves performance on par with supplying the actual paratope. We conclude that for docking simulations Parapred’s predictions are almost as informative as the actual paratope.

**Table 2.** The number of high, medium and low quality decoys obtained by running PatchDock with different constraints on a test set of 30 structures.

| Binding site | Top 10 |  |  | Top 200 |  |  |
| --- | --- | --- | --- | --- | --- | --- |
|  | *** | ** | * | *** | ** | * |
| CDRs | 0 | 2 | 0 | 1 | 14 | 0 |
| Paratope | 0 | 7 | 1 | 1 | 21 | 3 |
| Parapred | 0 | 8 | 0 | 1 | 19 | 2 |

We also measured the time taken by PatchDock to produce decoys on a machine with an “Intel(R) Core(TM) i7-6600U CPU @ 2.60GHz” processor (Table 3). We found that specifying Parapred’s predictions as a potential binding site produces a 1.7x speedup in PatchDock’s computations compared to specifying just the CDRs.

**Table 3.** PatchDock running time using different binding site constraints.

| Binding site | Running time |
| --- | --- |
| CDRs | 3h 50min 19.72s |
| Paratope | 2h 02min 22.50s |
| Parapred | 2h 11min 15.40s |
| **(1) vs (3) speedup:** | **1.75x** |

### C. Interpreting local neighbourhood features

The use of RNN allows our model to learn complex long-range dependencies between residues. This expressiveness also comes at a cost of interpretability—it is difficult to identify precisely which factors the model considers to be the most important. Nevertheless, we can inspect the model’s output and attempt to relate it to known findings to further confirm them or identify the model’s biases.

Figure 6 shows how frequently each residue type participates in binding, based on contact information from the dataset and our model’s predictions. Residue was considered to be binding if it had > 0.5 mean binding probability according to crossvalidation results. Overall, the model approximates the true binding profile well, showing particular preference for serine (S), tyrosine (Y), glycine (G) and threonine (T). Interestingly, the model largely ignores cysteine (C), most likely due to its tendency to form disulphide bonds and not therefore participate in binding.

**Fig. 6.**
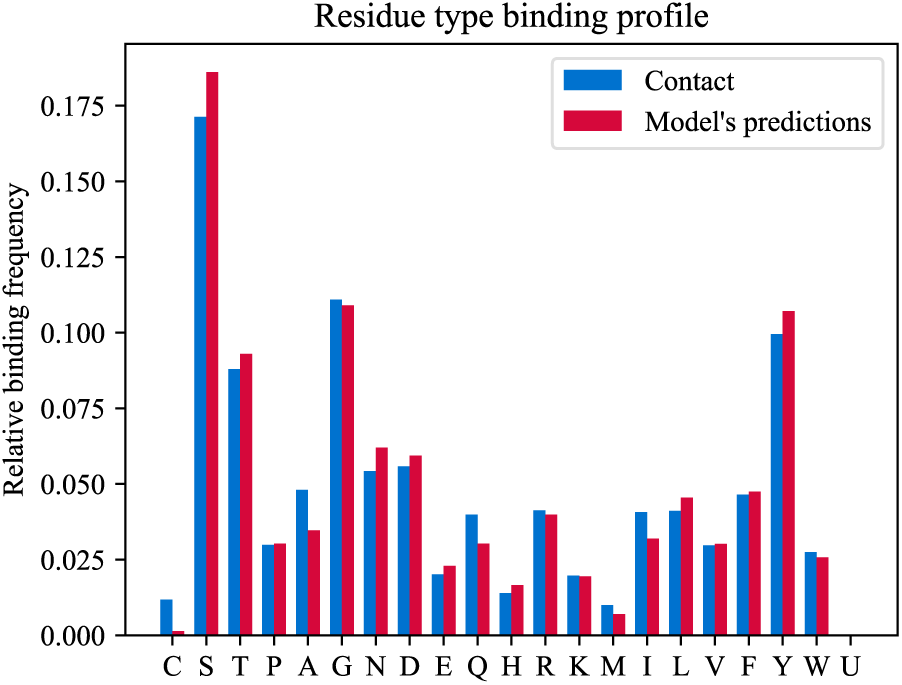
Frequency of residues of a particular type participating in binding, as computed from the dataset (blue) and as predicted by our model (red)

As discussed in the previous section, the first layer of our network consists 28 convolutional filters with kernel width 3, acting as local feature extractors. We can inspect what these filters learned by investigating which sequences of length 3 result in their highest response (*activation*). We measure activations after the the residual connection adds one-hot encoded residue type to the convolutional layer’s output; this allows to include the activation boost of each residue type for the first 20 filters due to one-hot encoding.

Table 4 shows neighbourhoods with highest activations for the first 20 filters (corresponding to each residue type). Findings include the following interesting points:

- Tryptophan (W) is present in many residue neighbour-• hoods, even though it does not often participate in binding itself.
- Kernel for lysine (K) does not actually consider lysine to be the most informative residue. It learns arginine’s (R) neighbourhoods.

**Table 4.**
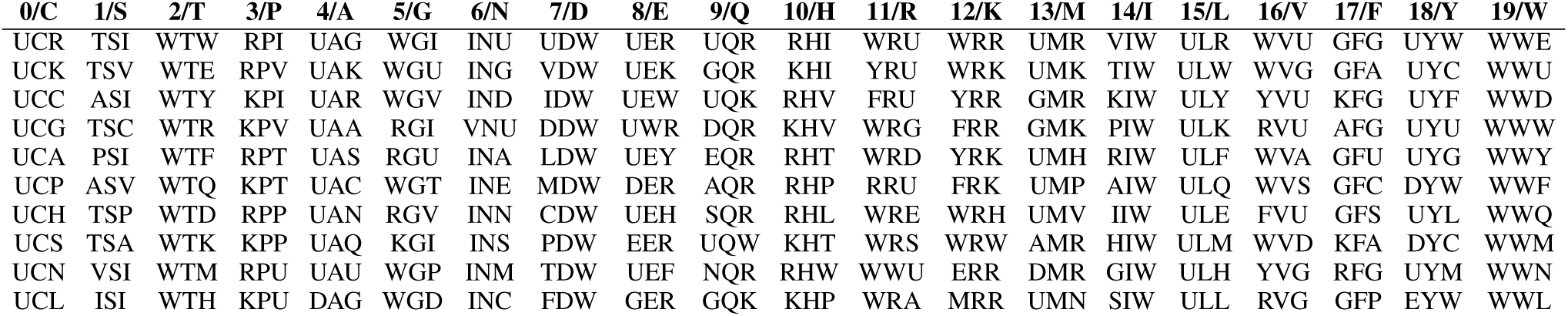
Top 10 sequences that activate the first 20 convolutional filters the most, sorted in descending order of the absolute value of activation. The presence of U indicates that the filter does not have a particular neighbour preference.

This investigation into model interpretability could be further extended by adding attention layers to the model (Bahdanau *et al.* [35]), which would allow inspecting positions in the input sequence that model learned to relate the most.

## IV. CONCLUSIONS

### A. Summary

To our best knowledge, this work is the first application of modern deep learning (CNN-and RNN-based neural networks) to the paratope prediction problem. Our model is able to generalise using only antibody sequence stretches corresponding to the CDRs (with 2 extra residues on the either side) and outperforms the current state-of-the-art by a statistically significant margin. We also showed that the model’s predictions provide speed and quality gains for the PatchDock rigid docking algorithm—decoy quality and time-to-dock were comparable to ones obtained when the docking algorithm has knowledge of the CDRs.

One of the main benefits of Parapred is that it does not rely on any higher-level antibody features: no full sequence, homology model, crystal structure or antigen information is required. However, if the user has such data already available, they might be able to usefully integrate it with the model, for example, by adding extra per-residue features.

## APPENDIX

*A. Extra features in the antibody residue encoding*

See Table 5.

**Table 5.** Extra features used in the amino acid (AA) residue encoding (U stands for the extra ‘unknown’ type): 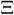—steric parameter, *α*_*p*_— polarisability, *v*_*v*_—volume, π—hydrophobicity, *I*—isoelectric point, *α*— helix probability, *β*—sheet probability.

| AA | $\Xi$ | $\alpha_p$ | $\nu_v$ | $\pi$ | $I$ | $\alpha$ | $\beta$ |
| --- | --- | --- | --- | --- | --- | --- | --- |
| C | 1.77 | 0.13 | 2.43 | 1.54 | 6.35 | 0.17 | 0.41 |
| S | 1.31 | 0.06 | 1.60 | -0.04 | 5.70 | 0.20 | 0.28 |
| T | 3.03 | 0.11 | 2.60 | 0.26 | 5.60 | 0.21 | 0.36 |
| P | 2.67 | 0.00 | 2.72 | 0.72 | 6.80 | 0.13 | 0.34 |
| A | 1.28 | 0.05 | 1.00 | 0.31 | 6.11 | 0.42 | 0.23 |
| G | 0.00 | 0.00 | 0.00 | 0.00 | 6.07 | 0.13 | 0.15 |
| N | 1.60 | 0.13 | 2.95 | -0.60 | 6.52 | 0.21 | 0.22 |
| D | 1.60 | 0.11 | 2.78 | -0.77 | 2.95 | 0.25 | 0.20 |
| E | 1.56 | 0.15 | 3.78 | -0.64 | 3.09 | 0.42 | 0.21 |
| Q | 1.56 | 0.18 | 3.95 | -0.22 | 5.65 | 0.36 | 0.25 |
| H | 2.99 | 0.23 | 4.66 | 0.13 | 7.69 | 0.27 | 0.30 |
| R | 2.34 | 0.29 | 6.13 | -1.01 | 10.74 | 0.36 | 0.25 |
| K | 1.89 | 0.22 | 4.77 | -0.99 | 9.99 | 0.32 | 0.27 |
| M | 2.35 | 0.22 | 4.43 | 1.23 | 5.71 | 0.38 | 0.32 |
| I | 4.19 | 0.19 | 4.00 | 1.80 | 6.04 | 0.30 | 0.45 |
| L | 2.59 | 0.19 | 4.00 | 1.70 | 6.04 | 0.39 | 0.31 |
| V | 3.67 | 0.14 | 3.00 | 1.22 | 6.02 | 0.27 | 0.49 |
| F | 2.94 | 0.29 | 5.89 | 1.79 | 5.67 | 0.30 | 0.38 |
| Y | 2.94 | 0.30 | 6.47 | 0.96 | 5.66 | 0.25 | 0.41 |
| W | 3.21 | 0.41 | 8.08 | 2.25 | 5.94 | 0.32 | 0.42 |
| U | 0.00 | 0.00 | 0.00 | 0.00 | 0.00 | 0.00 | 0.00 |

*B. Decoy classification*

Decoys are classified using the CAPRI criteria (Janin *et al.* [32], Méndez *et al.* [33]):

*ƒ*_NAT_: Two residues, one from the antibody and one from the antigen, are said to be an *interface pair* if they are sufficiently close to interact (here residues are assumed to be interacting if they are less than 5Å apart). *ƒ*_NAT_ is a proportion of interface pairs in the native complex that were reproduced in the generated one.

*L*_*RMS*_: The root mean square deviation (RMSD) of atoms’ coordinates in the generated orientation of the antigen compared to the true one (the lower the better). For two sets of coordinates u and v of size *n*, the RMSD is given by:

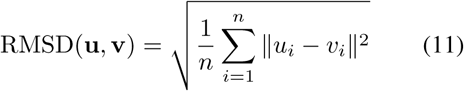

The RMSD is computed only for *backbone heavy atoms* (N, C, C_*α*_, O).

*I*_*RMS*_: The *L*_*RMS*_ measure unnecessarily penalises large antigen chains when the epitope has a roughly correct orientation but the rest of the chain does not. To better handle such cases, the *I*_*RMS*_ measure was introduced—the RMSD only of residues within 10Å of the epitope.

Decoys are assigned a class according to criteria in Table 6.

**Table 6.** Decoy classification using *ƒ*_NAT_, *L*_*RMS*_ and *I*_*RMS*_ measures. The decoy has to match the *ƒ*_NAT_ criteria and either the *L*_*RMS*_ or *I*_*RMS*_ criteria.

| Class | $f_{\text{NAT}}$ | $L_{RMS}$ | $I_{RMS}$ |
| --- | --- | --- | --- |
| High | $\geq 0.5$ | $\leq 1.0$ | or $\leq 1.0$ |
| Medium | $\geq 0.3$ | $1.0 < \cdot \leq 5.0$ | or $1.0 < \cdot \leq 2.0$ |
| Acceptable | $\geq 0.1$ | $5.0 < \cdot \leq 10.0$ | or $2.0 < \cdot \leq 4.0$ |
| Incorrect | $< 0.1$ | — | — |

*C. Dataset*

See Table 7.

**Table 7.**
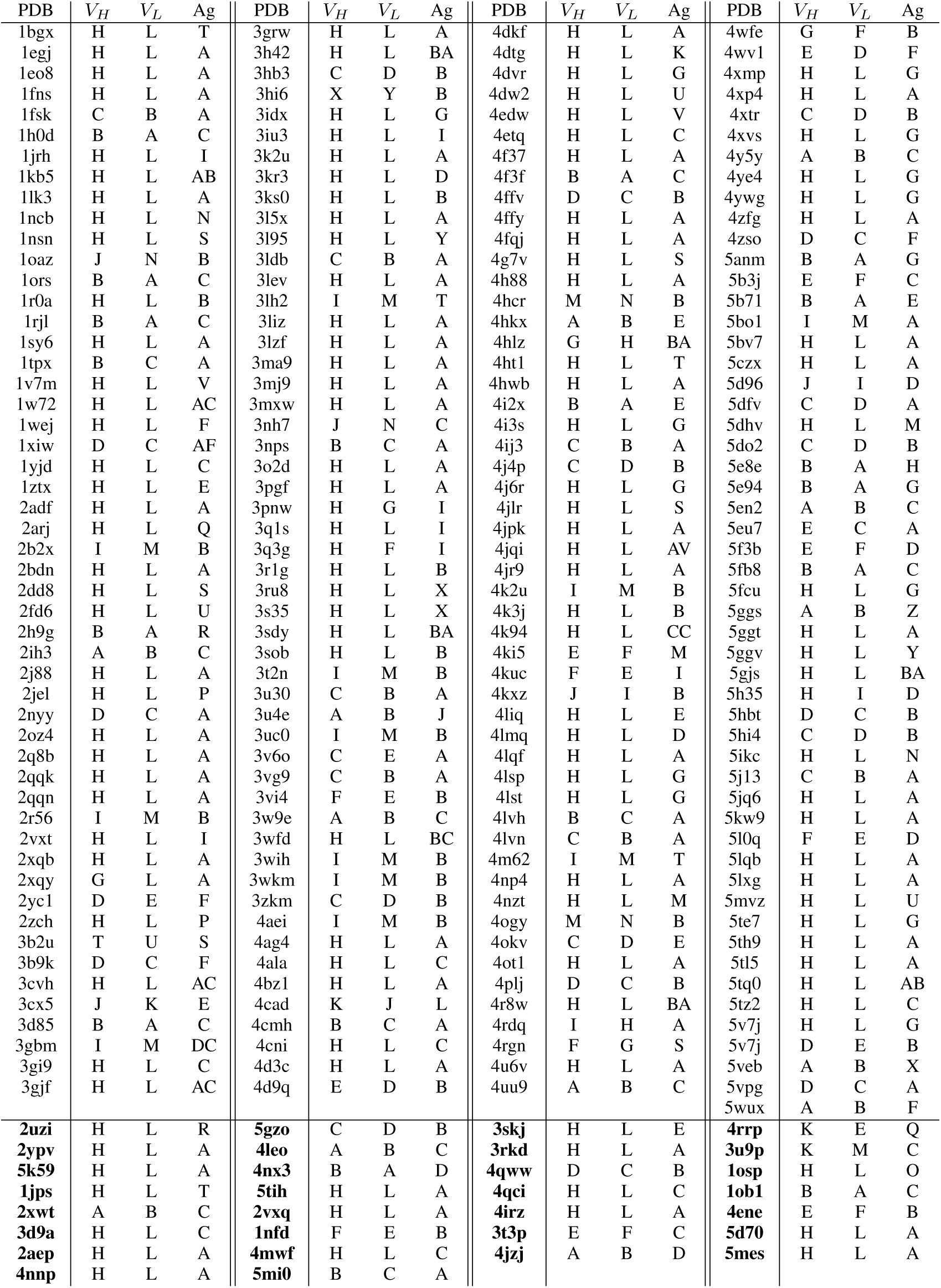
239 bound antibody-antigen complex dataset used to train our model. Each entry shows the PDB code, as well as antibody heavy, antibody light and antigen chain IDs. Structures in bold were used as a test set for the docking experiment.

